# Exposure during embryonic development to Roundup® Power 2.0 affects lateralization, level of activity and growth, but not defensive behaviour of marsh frog tadpoles

**DOI:** 10.1101/847251

**Authors:** Alessandro Bolis, Andrea Gazzola, Daniele Pellitteri-Rosa, Anita Colombo, Patrizia Bonfanti, Adriana Bellati

**Author notes:** <u>corresponding author:</u> Adriana Bellati, Department of Earth and Environmental Sciences, University of Pavia, Via Ferrata 9, 27100 Pavia, Italy.

## Abstract

As glyphosate-based herbicides, sold under the commercial name Roundup®, represent the most used herbicides in the world, contamination of the freshwater environment by glyphosate has become a widespread issue. In Italy, glyphosate was detected in half of the surface waters monitoring sites and its concentrations were higher than environmental quality standards in 24.5% of them. It can last from days to months in water, leading to exposure for aquatic organisms and specifically to amphibians’ larvae that develop in shallow water bodies with proven effects to development and behaviour. In this study, we tested the effects of a 96h exposure during embryonic development of marsh frog’s tadpoles to three ecologically relevant Roundup® Power 2.0 concentrations. As expected, given the low concentrations tested, no mortality was observed. Morphological measurements highlighted a reduction in the total length in tadpoles exposed to 7.6 mg a.e./L, while an increase was observed at lower concentrations of 0.7 and 3.1 mg a.e./L compared to control group. Tadpoles raised in 7.6 mg a.e./L also showed a smaller tail membrane than those raised in the control solution. Regarding behaviour, we tested tadpoles in two different sessions (Gosner stages 25 and 28/29) for lateralization, antipredator response and basal activity. Lower intensity of lateralization was detected in tadpoles raised at the highest Roundup® concentration in the first session of observation, while no significant difference among treatments was observed in the second one. In both sessions, effects of glyphosate embryonic exposure on antipredator response, measured as the proportional change in activity after the injection of tadpole-fed predator (*Anax imperator*) cue, were not detected. Tadpoles exposed during embryonic development to Roundup® exhibited lower basal activity than the control group, with the strongest reduction for the 7.6 mg a.e./L treatment. Our results reinforce the concern of glyphosate contamination impact on amphibians.

## INTRODUCTION

Glyphosate-based herbicides (GBHs) are between the most sold broad-spectrum herbicides in the world (Duke and Powles, 2008), whose use reaches nearly 100 million kilograms on an annual basis (Grube et al., 2011). The active ingredient, glyphosate, appears in several alternative formulations, altogether known under the commercial name Roundup® (Monsanto Co., St. Louis, MO, USA). Glyphosate use spans different applications, from no-tillage farming and conventional agriculture to non-cultivated areas, forest management and private gardening (Dill, 2005; Dill et al., 2010). Its use is approved until 2022 by the European Union (European Commission, 2017), however in Italy its application is restricted during harvest and threshing, and banned in public gardens and parks, schools, private gardening and health facilities (Italian Ministry of Health, 2016). Although the aforementioned restrictions, a recent report concerning surface waters pesticide contamination highlighted the presence of this molecule in 47.4% of the Surface Waters Monitoring Sites in Italy (SWMS), with its concentration exceeding the Environmental Quality Standards (EQS) in 24.5% of them (ISPRA, 2018).

As this molecule has been engineered to kill plants, specifically post-emergent leaves and grasses, glyphosate is considered to have low mammalian toxicity, being “practically non-toxic” for bees, birds and most aquatic organisms, according to the World Health Organization/Food and Agriculture Organization (Lajmanovich et al., 2011). Microorganism degrade the active ingredient into its metabolite aminomethylphosphonic acid (AMPA), which is eventually oxidized into carbon dioxide, in both soil and water. Since the rate of degradation is strictly dependent on physical (e.g. temperature) and chemical (e.g. pH) environmental conditions, glyphosate’s half-life can last from days to months (e.g. 7–70 days in the water; Giesy et al., 2000), leading to possible chronic exposures for organisms living in certain environments (Feng et al., 1990; Borggaard and Gimsing, 2008; Bailey et al., 2018). GBH formulations are generally more toxic than glyphosate by itself, mainly due to the presence of surfactants (e.g. polyoxyethylene - POE, polyethoxylated tallow amine - POEA). Previous studies showed that the toxicity of GBHs to aquatic organisms is largely caused by the surfactant in the mixture (Edginton et al., 2004; Moore et al., 2012). These substances may not only cause toxicity by themselves, but also facilitate the penetration of the active ingredient in animal cells causing teratogenic effects, as shown in amphibian embryos and tadpoles of several species (Borggaard and Gimsing, 2008; Paganelli et al., 2010; Bonfanti et al., 2018; Gill et al., 2018).

To date, GBHs have been proved to affect amphibians’ development through growth retardation and weight reduction during larval stages, especially in anurans (Smith, 2001; Cauble and Wagner, 2005; Lanctôt et al., 2014; Navarro-Martín et al., 2014). Furthermore, GBHs have been shown to alter behaviour in aquatic organisms, like fishes and amphibians, involving locomotion, memory, visual and olfactory senses and antipredator responses (Tierney et al., 2006; Moore et al., 2015; Besson et al., 2017; Bridi et al., 2017; Mikò et al., 2017). In the poecilid *Cnesterodon decemmaculatus*, exposure to GBHs during development has been proven to cause a significant inhibitory effect on AChE (acetylcholinesterase) activity in the tail (Menéndez-Helman et al., 2012). The inhibition of this enzyme, involved in the breakdown of the neurotransmitter acetylcholine in muscle and nerve synapses and thus in the end of a transmission of a neural impulse (Zimmerman and Soreq, 2006; Tripathi and Srivastava, 2010), leads to a possible reduction of basal activity. Nonetheless, another study demonstrated that AChE activity increases in tadpoles of the marsh frog *Pelophylax ridibundus*, the green toad *Bufotes viridis*, and the African clawed toad *Xenopus laevis* exposed to GBH formulations (Güngördü, 2013). This finding however is still unmatched by other studies, at least for anurans since glyphosate instead decreased AChE activity in tadpoles of the toad *Rhinella arenarum* in a study by Lajmanovich and colleagues (2011).

An additional negative effect of herbicide water contamination could be lateralization impairment (Besson et al., 2017). Brain lateralization is the difference between the activity of the right and left hemisphere in the execution of several tasks, implying the preferential use of one body portion over the other (Davidson and Hugdahl, 1996; Bisazza et al., 1998). Laterality is thought to augment cognitive abilities, by optimizing the processing of information in the two-separate brain hemispheres, thus enhancing the ability to make decisions when facing novel multi-sensory stimuli (Vallortigara and Rogers, 2005; Salva et al., 2012). Defence against predators is a well-known lateralized behaviour in vertebrates, as the recognition and escape performances seem to depend on the side of appearance of a predatory threat (Siniscalchi et al., 2010; Shibasaki et al., 2014). Fishes, birds, and reptiles have shown eye preference when keeping track of predators or choosing for a prevalent escape direction in predatory risk conditions (Sovrano et al., 2005; Koboroff et al., 2008; Martín et al., 2010; Pellitteri-Rosa and Gazzola, 2018).

In larval amphibians, defensive behaviour is generally activated by chemical (olfactory) cues, released in the water both by injured conspecifics (alarm cue) as well as by predators (kairomones). It has been shown that glyphosate negatively affects the functionality of the olfactory system by inactivating the chemical cues conveying information on predation risk, thus inhibiting predator recognition, or by lowering the learning ability of tadpoles (Mandrillon and Saglio, 2007a, b; Moore et al., 2015). An altered olfactory system could in turn weaken defensive responses and negatively affect tadpoles’ survival in their natural environment.

Behavioural alterations may appear when tadpoles develop in the contaminated medium (i.e. chronic exposure), or alternatively when adults are exposed, even briefly, to the substance (i.e. acute-transitory exposure). Therefore, contaminants may affect amphibians at every life stage, with negative effect on growth and reproduction, and finally survival of the single individual likewise the entire population (Relyea, 2005; Gill et al., 2018). The complex life cycle of anurans, which shifts their biological cycle from water to terrestrial habitats and the assumption that several environmental cues experienced during embryonic development can affect the range of physiological and behavioural responses available as adults (Dufty et al., 2002), makes them an ideal candidate to explore the effects of pollutants during development.

In this study, we explored the effects of glyphosate on the morphology and some fitness-related behavioural traits in tadpoles of a widely distributed European anuran, the marsh frog *P. ridibundus*, by exposing embryos to different ecologically relevant concentrations of the commercial GBH Roundup® Power 2.0 (Monsanto Co., St. Louis, MO, USA). *P. ridibundus* eggs are frequently found in areas with vegetation, wetlands, agricultural areas, and urban regions where the presence of pesticides could be significant. Our aims were to investigate the possible effects of early (embryonic) exposure to RU-PW on: (1) total length and tail depth, to detect growth retardation; (2) lateralization, as a proxy of the correct development of the nervous system of the tadpole; (3) level of activity (both basal and in the presence of a predatory stimulus), to infer the ability of the individual to correctly cope with important environmental pressures (i.e. predation). Previous studies showed that *P. ridibundus* tadpoles are more resistant to glyphosate-based herbicide than other species of anurans (Güngördü, 2013). Nonetheless, adults of this species demonstrated a slight increase in the hepatosomatic index (HSI) when exposed via intraperitoneal to 0.138 × 10^−3^ mL Roundup®/g per body mass (Păunescu and Ponepal, 2011). According to previous findings, we expect to find a relationship between glyphosate exposure and both tadpoles morphological and behavioural traits. At the same time, we also expect these effects to only appear at particular developmental stages and have little or no effect on the long-term survival of individuals, as a result of general high resistance of the species to environmental stressors, like xenobiotics.

## MATERIALS AND METHODS

### Animals collection and breeding

The marsh frog is an anuran endemic to Central and Eastern Europe. In Italy, native populations of this taxon only occur at the edge of the Northern-Eastern region of Friuli-Venezia Giulia (Lanza et al., 2006). Nonetheless, several populations of this frog had been introduced in Italy for edible purposes. According to the Environmental Impact Classification of Alien Taxa (EICAT) of the International Union for Conservation of Nature (IUCN), the species is considered one of the invasive species with the highest potential to impact native species diversity worldwide, because of its ability to occupy a wide variety of habitats on one hand, and to hybridize with native taxa, thus producing viable and fertile hybrids, on the other (Kumschick et al., 2017).

Adult marsh frogs were collected at night on June 15^th^, 2018, from an artificial pond situated in the municipality of Mezzolombardo (46.19°N, 11.09°E, TN, Italy). Here, previous surveys had highlighted the presence of water frogs that were ascribed to *P. ridibundus* following molecular analysis. Frogs were captured using fishing nets and transported to the laboratory of the Department of Earth and Environmental Sciences of the University of Pavia. Until the beginning of the 96h exposure, frogs were kept in plastic tanks filled with dechlorinated tap water and fed with crickets (*Acheta domesticus*) ad libitum.

Embryos of *P. ridibundus* used in the 96h exposure were obtained through in vitro fertilization following the procedure for anuran breeding provided by Pruvost (2013), and Berger et al. (1994) with modifications (Bonfanti et al., 2004). Specifically, 24 h before egg collection, we triggered ovulation stimulation by injecting both females and males (n = 6) with LHRH hormone (BachemH-7525, Bubendorf, Switzerland) at a concentration of 2 mg in 100 mL Holtfreter’s solution (Holtfreter, 1944). Individuals were injected into the dorsal lymph sac 10 µl/g of body mass of hormone.

The male selected for the crossing was euthanized in a MS-222 solution (Sigma A-5040, St. Gallen, Switzerland) at 2 mg/L. Testes were removed, put in 2 mL of cold DBT solution (Tris-HCL 10 m, pH 7.5) and minced to obtain a sperm suspension. Egg groups, obtained by gently massaging the abdominal region of the females, were collected in 90-mm plastic Petri dishes and immediately inseminated with sperm suspension; after 2 min, 30 ml of Holtfreter’s solution were added to each Petri dish. Successful insemination was detected when after 30 min of incubation at 25 °C all the eggs were oriented with the dark side (animal pole) up. Embryos (Gosner stages 4-5) were selected under a stereomicroscope and the jelly coat was removed by swirling the embryos for 1-2 min in a 2.25% L cysteine solution (pH 8.1). All the fertilizations were performed on June 18^th^, 2018.

We collected five dragonfly larvae (*Anax imperator*) from a small pond situated in the botanical garden in Pavia (45.18°N, 9.15°E, Italy) on July 2^nd^, 2018. They were individually kept in 800 mL cups filled with 500 mL of aged tap water and wooden sticks as perching sites. These animals were later used for behavioural trials.

### Roundup® Power 2.0 solutions

Roundup® Power 2.0 (Monsanto Italia S.P.A.), referred to as RU-PW, was formulated with a guarantee of 360 g glyphosate acid equivalent (a.e.) per litre present as the potassium salt (CAS RN 70901-12-1), six percent by volume of ethoxylated ether alkyl ammine (CAS RN 68478-96-6) and 58.5% water and other ingredients not specified by the producer.

A stock solution of RU-PW at nominal concentration of 100 mg a.e./L using FETAX solution was prepared. In according to previous experiments, we opted to use FETAX solution as control, since it is the optimal solution for the development of water frogs. The composition in mg/L was: NaCl 625, NaHCO_3_ 96, KCl 30, CaCl_2_ 15, CaSO_4_-2H_2_O 60, and MgSO_4_ 70, pH 7.5–8.5 (Dawson and Bantle, 1987).

### 96h exposure test

Normally-cleaved embryos at the midblastula stage (Gosner stage 8; Gosner, 1960) five hours post fertilization (hpf), were selected from each female (n = 2), assigned to the experimental groups (n = 15 per group) and placed in covered Petri dishes containing 40 mL of control (FETAX) or RU-PW solution. For each treatment (three RU-PW concentrations and control) and for each female, five replicas were performed for a total of 300 embryos per female.

RU-PW concentrations used for the 96h exposure were 0.7 mg a.e./L, 3.1 mg a.e./L and 7.6 mg a.e./L.; control treatment was represented by FETAX solution. Each experimental concentration corresponded to one of the following three different scenarios: environmental concentration of the herbicide without intervention (0.7 mg a.e./L), concentration occurring shortly after the application of the herbicide (3.1 mg a.e./L), and concentration estimated in worst-case scenarios (e.g. direct spraying in a flooded field, 7.6 mg a.e./L) (Wagner, 2013). Embryos were incubated in a thermostatic chamber at 25 ± 0.5 °C and exposure solutions were renewed every 24 h (semi-static conditions). The selected concentrations should not cause different mortality rates with respect to the control group, since the estimated LC50 for RU-PW is 24.75 mg a.e./L for *Xenopus laevis* (Bonfanti et al., 2018), a species considered more sensitive to Roundup than marsh frogs (Güngordü, 2013).

### Morphological measurements

At the end of the exposure period (96 hpf) we randomly selected a total of 220 tadpoles equally balanced both for treatment and female. They were euthanized and formalin fixed and photographed through a stereomicroscope equipped with a camera (AxioCam ERc5s) to estimate growth retardation by measuring the total length and maximum tail height (Altig, 2017), using the digitizing software AxioVision. Remaining individuals were reared in 16 plastic tanks (2 tanks per treatment and female) filled with dechlorinated tap water (8 L), under natural light conditions. After hatching, tadpoles were fed with rabbit food, and reunited in four plastic tanks according to treatment (control, 0.7 mg a.e./L, 3.1 mg a.e./L, 7.60 mg a.e./L solutions) on July 2^nd^.

### Behavioural trials: rotational preference, antipredator response and basal activity

We began the behavioural trials when tadpoles reached Gosner developmental stage 25 on July 3^rd^. We recorded the activity of 200 tadpoles (n = 50 from each treatment) in the span of three days, from 9:00 am to 4:00 pm. The tests consisted in recording the activity of individuals for 15 min, while measuring three distinct behavioural variables: rotational preference (i.e. lateralization), antipredator response and basal activity.

Before starting the trials, each individual was placed in 100 × 20 mm Petri dishes filled with 60 ml of well-aged tap water for a period of acclimation (15 min). To measure rotational preference, we recorded the time spent swimming in both clockwise and counterclockwise direction during the first 10 min of the test. This is a well-established method for assessing lateralization in tadpoles (Vandenberg and Levin, 2013; Lucon-Xiccato et al., 2016). Only clockwise and counterclockwise movements made in the outer portion of the Petri dishes were measured. We recorded lateralization only when tadpoles’ distance from the centre of the Petri dishes was at least 3 cm. Tadpoles that did not move for the whole test were excluded from the analysis (12 tadpoles in the first session and no tadpoles in the second one).

Tadpoles of many frog species are known to reduce level of activity after being exposed to predatory cues, in particular when chemicals are produced by a familiar predator preying upon conspecific prey (Ferrari and Chivers, 2009). Accordingly, for each trial we collected 1 mL of water from four different plastic tanks (500 ml) containing *Anax imperator* larvae, which were fed on conspecific tadpoles (one per dragonfly larvae) at least one hour before the beginning of the recording sessions. This procedure allowed us to have freshly produced odours, and to use them as a reliable olfactory signal consisting in predatory kairomones released by predators, and tadpoles’ alarm cue (Hettyey et al., 2015).

The antipredator response was estimated by recording the amount of time (in seconds) the tadpoles were active in a five minutes time span both before and after the injection of the stimulus, which consisted in 1 mL of either water (blank) or alarm cue. In order to detect possible differences within tadpoles from different developmental treatment in antipredator responses, the movement after and before the injection were compared. We considered the activity recorded before the injection of water or alarm cue, measured in a 5 min period, as the basal activity level.

Three weeks later, when tadpoles reached Gosner stage 28/29, we repeated the experiment using the same experimental protocol as explained above and recorded the activity of 80 tadpoles (n = 20, from each developmental treatment).

### Statistical analysis

We applied the non-parametric test Kruskall-Wallis to investigate differences in the total length, and Dunn’s test of multiple comparisons with p-values adjusted with the Benjamini-Hochberg method to compare control with other treatments. We then compared maximum tail height in tadpoles exposed to RU-PW to those raised in the control solution with a robust bootstrap version of ANCOVA for trimmed means with the total length as covariate (bootstrap = 10,000, 20% trimmed means) using the R package WRS2, function: *ancboot* (Wilcox and Mair, 2016). This test compares trimmed means at different points along the covariate and finds five points where the slopes are roughly the same, then it compares the trimmed means at these points and explore the possible differences. Since this analysis can be performed only when comparing two groups, the control group was compared to the three RU-PW treatments in three different analysis.

Rotational preference was analysed through two parameters often used to study lateralization (Cantalupo et al., 1995). Lateralization directionality (L_R_ index) was calculated with the following formula: (clockwise swimming time – counterclockwise swimming time) / (clockwise swimming time + counterclockwise swimming time) × 100. Moreover, the intensity of lateralization (L_A_ index) equals to the absolute value of L_R_ (L_A_ = |L_R_|). We compared both indexes between tadpoles from different developmental treatments using a bootstrap version of one-way ANOVA for trimmed means (R package: WRS, function: *t1waybt*), both for the first and second session (bootstrap = 10,000, 20% trimmed means; Wilcox, 2011). We then compared indexes between the two experimental sessions with a two-way ANOVA for trimmed means (R package: WRS2, function: *t2way*). Post-hoc tests were executed with *lincomb* function (R package: WRS).

Regarding the antipredator response, we calculated change in activity using the following formula: (movement after – movement before) / (movement before) the injection of the stimulus. In order to explore the disturbance effect of the injection procedure we used a non-parametric Wilcoxon rank sum test to compare the tadpoles mean level of activity before and after the water injection for each treatment in both experimental sessions. A two-way ANOVA for trimmed means was used to compare change in activity between groups and experimental session (R package: WRS2, function: *t2way*). For each session, we used one-way ANOVA for trimmed means to explore differences among embryonic treatments for the level of activity.

Finally, we used a linear model (*lm*) to explore differences in the basal activity level and included embryonic treatment, experimental session and their interaction as main predictors. Planned comparisons with control group in each session were extracted from the model with “emmeans” R-package and no *p*-value correction (Lenth, 2018).

## RESULTS

### Morphology

We detected significant differences between treatments regarding the measured morphological parameters (Fig. 1). The total length was different between embryonic treatments (X^2^ = 50.0, *P* < 0.001). Tadpoles exposed to low concentrations RU-PW (0.7 and 3.1 mg a.e./L) showed a higher total length than those raised in the control solution (Z = −3.01, *P* = 0.003; Z = −3.27, *P* = 0.002 respectively). On the contrary, tadpoles which developed in the highest concentration (7.6 mg a.e./L) showed a lower total length compared to the control group (Z = 2.26, *P* = 0.008). Results from robust bootstrap version of ANCOVA for trimmed means showed that significative differences in the maximum tail height emerged only for the highest glyphosate treatment (7.6 mg a.e./L) in comparison to the control group (Table 1).

**Figure 1:**
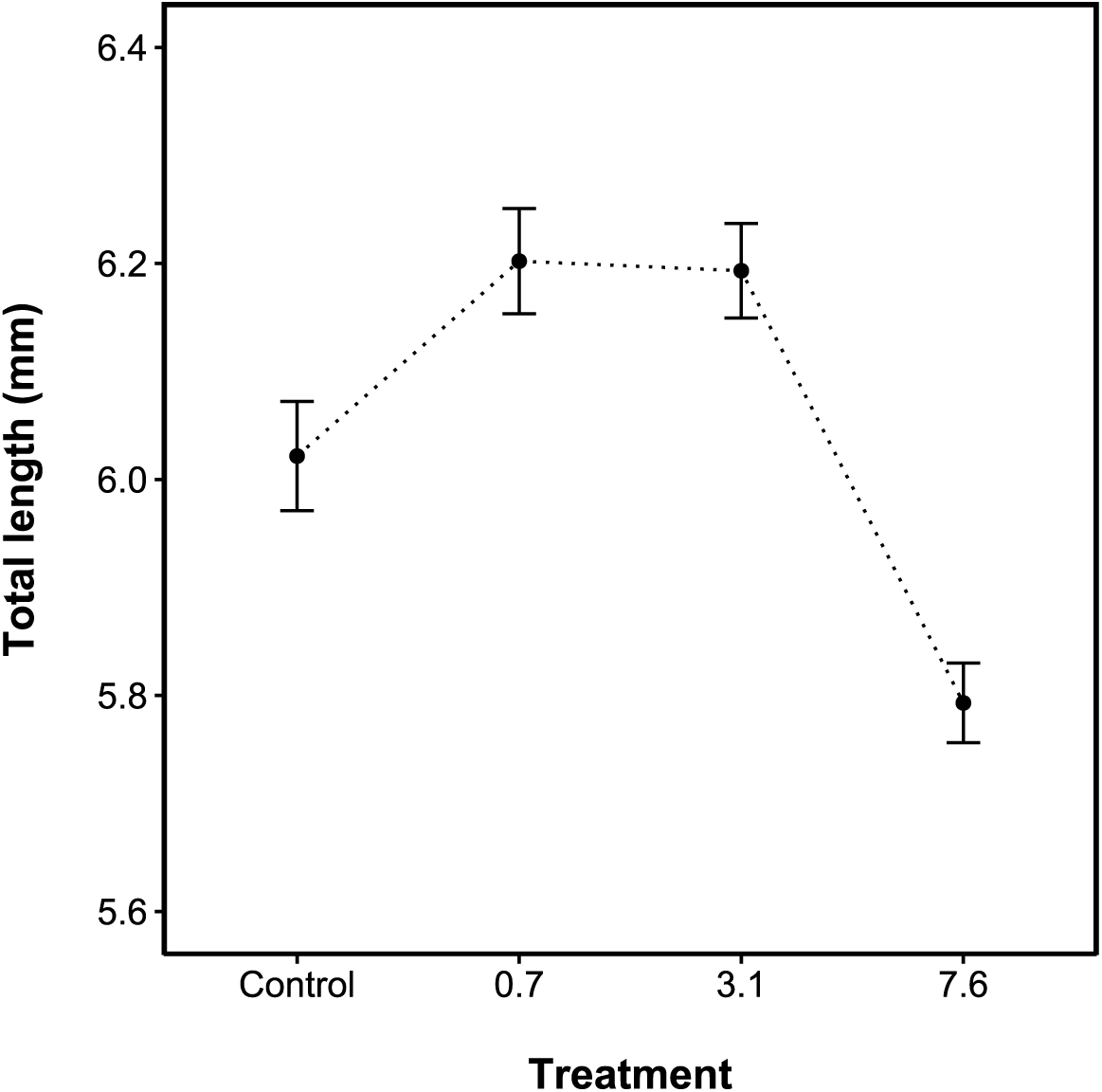
Mean ± S.E. of total length in the four treatment groups. Dashed line highlights the hormetic effect of RU-PW on the growth rate. Concentrations are reported in mg a.e./L.

**Table 1:** Results from Robust ANCOVA analysis (20% trimmed means, bootstrap = 10.000). The *design points* are total length (covariate) values (µm) for which the relationship between total length and maximum tail height are comparable in the compared groups. *N1* and *N2* are the number of individuals that have a covariate value close to the *design points*. The *difference* column shows the difference in the trimmed means for the maximum tail height between the two groups and the test statistic values are stored in the *F* column (*difference/*S.E.) with the corresponding *p*-values in the *P* column. 95% confidence intervals are in the *LCI95%* (lower) and *UCI95%* (upper).

|  | *design points* | *N1* | *N2* | *difference* | *LCI 95%* | *UCI 95%* | *F* | *P* |
| --- | --- | --- | --- | --- | --- | --- | --- | --- |
| ***C – T1*** | 5612.60 | 32 | 14 | -53.97 | -127.37 | 19.43 | -2.17 | < 0.05 |
|  | 5736.65 | 36 | 22 | -58.16 | -128.33 | 12.00 | -2.44 | < 0.05 |
|  | 5958.68 | 40 | 33 | -59.99 | -136.11 | 16.12 | -2.32 | < 0.05 |
|  | 6221.33 | 28 | 42 | -25.74 | -107.76 | 56.28 | -0.93 | 0.37 |
|  | 6734.02 | 13 | 16 | -3.42 | -104.86 | 98.02 | -0.09 | 0.92 |
| ***C – T2*** | 5690.44 | 33 | 12 | -86.17 | -149.54 | -22.80 | -3.86 | <0.001 |
|  | 5854.84 | 42 | 23 | -54.98 | -115.72 | 5.75 | -2.57 | < 0.05 |
|  | 6018.72 | 40 | 35 | -43.00 | -114.78 | 28.77 | -1.70 | 0.10 |
|  | 6265.36 | 27 | 35 | 12.17 | -69.08 | 93.43 | 0.43 | 0.67 |
|  | 6691.35 | 13 | 13 | 11.71 | -92.27 | 115.68 | 0.32 | 0.75 |
| ***C – T3*** | 5512.82 | 27 | 20 | 272.15 | 200.15 | 345.05 | 10.58 | < **0.001** |
|  | 5631.46 | 33 | 32 | 283.85 | 223.67 | 344.03 | 13.27 | < **0.001** |
|  | 5735.77 | 36 | 36 | 287.59 | 225.15 | 350.03 | 12.95 | < **0.001** |
|  | 5894.17 | 43 | 25 | 301.28 | 234.04 | 368.51 | 12.61 | < **0.001** |
|  | 6018.72 | 40 | 17 | 303.48 | 233.18 | 373.78 | 12.14 | < **0.001** |

### Lateralization

In the first behavioural session, carried out at Gosner stage 25, L_A_ index was not equal among developmental treatments (F = 3.34, *P* = 0.024, d = 0.28; Fig. 3). The L_A_ index was significantly higher in the control group compared to tadpoles raised in the highest treatment concentration tested (7.6 mg a.e./L; Ψ= 22.22, *P* = 0.003), however no significant difference was detected between control and both 0.7 and 3.1 mg a.e/L (lower *p*-value: Ψ = 13.25, *P =* 0.07). In the second session (Gosner stage 28/29), statistically significant differences among treatments were no longer observed (F = 1.66, *P* = 0.20, d = 0.33). The global analysis for trimmed means showed a significant effect for the embryonic treatment (F = 10.48, *P* = 0.02) and for the experimental session (F = 62.32, *P* = 0.001), but not for the interaction of the two (F = 4.76, *P* = 0.20; Fig. 2). In all four groups L_A_ index values decreased from the first to the second session (Fig. 2).

**Figure 2:**
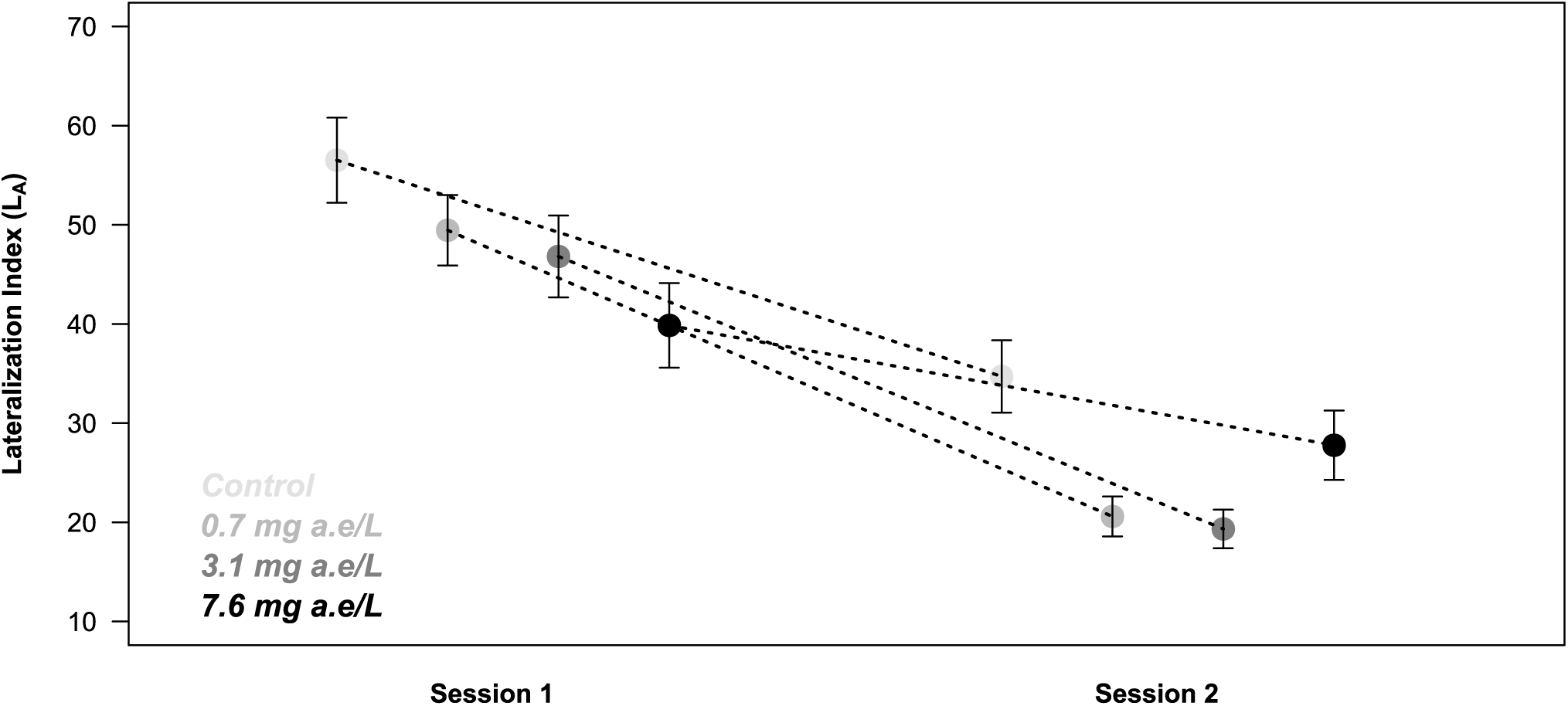
LA index for all groups in the first and second trial session. Values are mean ± S.E.

**Figure 3:**
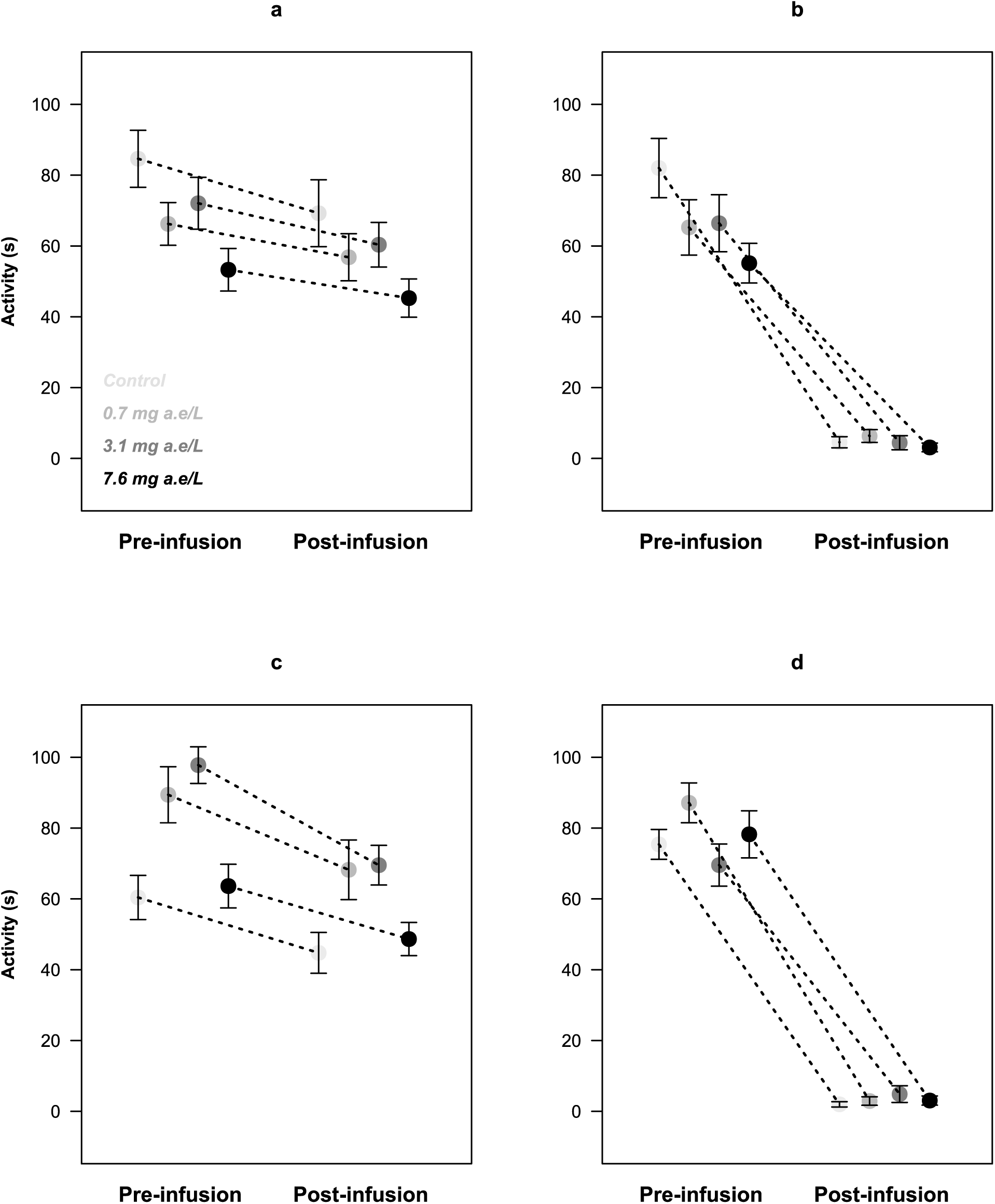
Activity before and after injection of water (**a**, **c**), predatory and conspecific cue (**b**, **d**) in the first (**a**, **b**) and the second session (**c**, **d**).

Tadpoles did not exhibit directional preference for L_R_ index, and we did not observe differences between the treatment groups in both sessions (Session 1: F = 0.46, *P =* 0.69, d = 0.11; Session 2: F = 0.27, *P* = 0.83, d = 0.15).

### Antipredator response

In both sessions (Gosner stages 25 and 28/29), the level of activity before and after the injection of water stimulus was not significantly different for all treatments (highest difference: W = 210, *P =* 0.11; Fig. 3a and 3c), thus showing a negligible disturbance effect of the injection procedure. The sole exception was represented by 3.1 mg a.e./L treatment in the second session, which significantly reduced activity level after the water stimulus injection (W = 21, *P* = 0.02; Fig. 3c). Concerning the antipredator response, we did not detect any significant difference in tadpoles’ activity change among embryonic treatments after the injection of the stimulus (either water or predatory cue; *P* = 0.75; Fig. 3b and 3d). Nevertheless, chemical stimulus and experimental session seemed to have a significant effect for the proportional change in the level of activity (value= 1087.24, *P* < 0.001 and value=5.36, *P* = 0.02, respectively). All groups showed a strong significant decrease in the level of activity after the injection of predatory cue in comparison to water injection (Session 1: F = 979.22, *P <* 0.001; Session 2: F = 1721.90, *P* < 0.001). No significant interaction was detected between embryonic treatment and the type of stimulus injected (Session 1: F = 1.45, *P =* 0.70; Session 2: F = 2.43, *P* = 0.53).

### Basal activity

The global linear model highlighted a significant interaction between embryonic treatment and experimental session (F = 3.23, *P* = 0.02; Fig. 4). In the first session (Gosner stage 25) we observed that basal activity, measured in a 5 min time span, was significantly higher in tadpoles raised in the control solution than in those exposed to the herbicide (F = 5.62, *P* = 0.001). In the first session, the intermediate RU-PW concentration groups (0.7 and 3.1 mg a.e./L) showed a significant reduction in the basal activity compared to the control (t.ratio = −2.54, *P* = 0.01 and t.ratio = −2.04, *P* = 0.04, respectively). Tadpoles exposed to the highest concentration of the herbicide (7.6 mg a.e./L) exhibited the strongest reduction in the basal level of activity (t.ratio = − 4.23, *P* < 0.001) when matched to the control group. In the second session (Gosner stage 28/29), tadpoles raised with RU-PW were overall more active than the control group but no significant difference was detected (highest difference: t.ratio = −1.79, *P*= 0.07).

**Figure 4:**
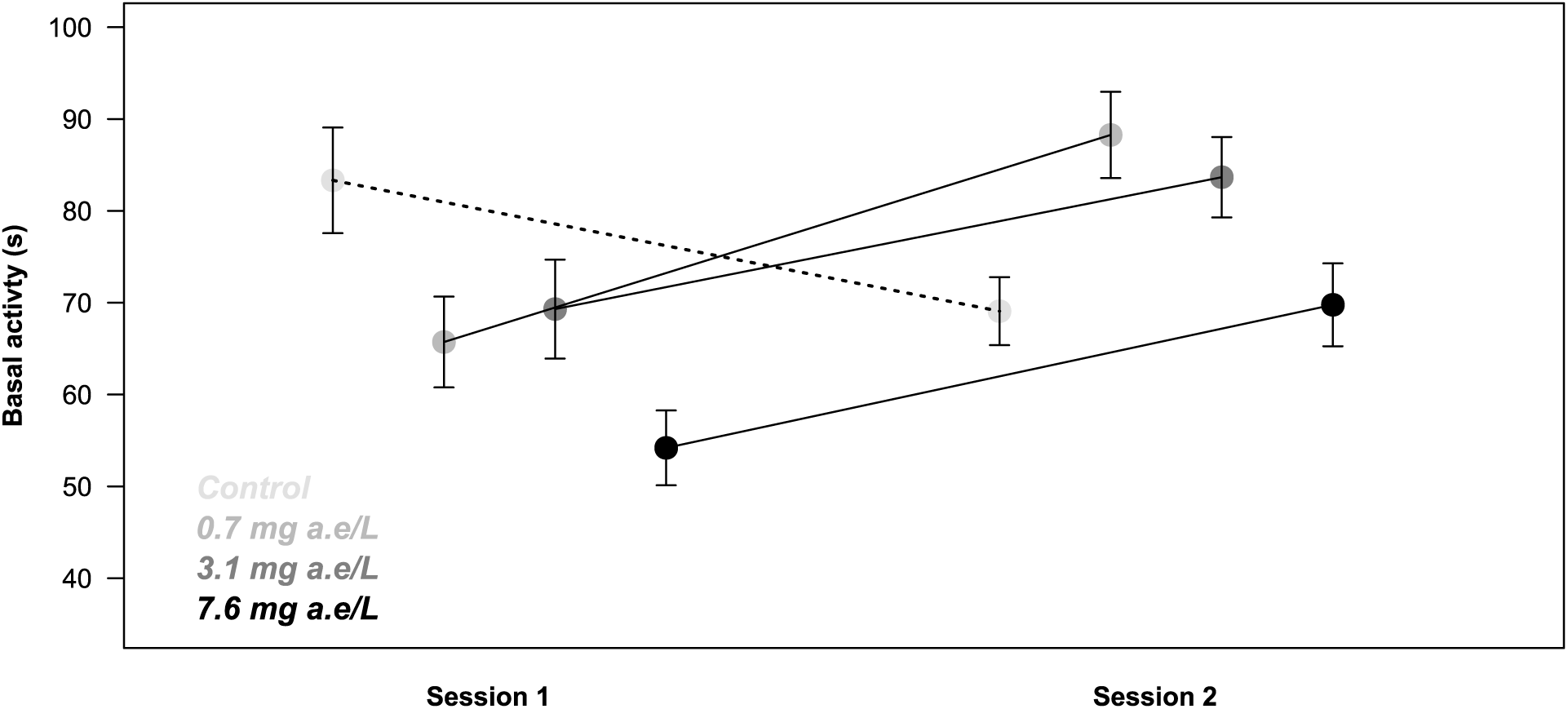
Basal activity levels (in seconds) for both sessions. Values are mean ± S.E.

## DISCUSSION

Our study clearly supports the hypothesis that GBHs in freshwater habitat could affect the life cycle of marsh frog’s tadpoles in terms of development, lateralization and activity level, all factors closely related to the ability of the larvae to cope with environmental pressures. As a general consideration, the absence of mortality rate in the different groups tested (both control and exposed) indicates a good quality of the experimental conditions and confirms the sub-lethal concentrations of GBHs. Similarly, no mortality has been observed in experiments using tadpoles exposed to ecologically relevant concentrations of several herbicides (Dornelles and Oliveira, 2014; Wilkens et al., 2019).

### Effects of RU-PW on morphological traits

Analysis of the morphological parameters revealed that growth of marsh frog tadpoles was affected by ecologically relevant glyphosate concentrations after the 96h laboratory exposure. Tadpoles exposed to low and intermediate concentrations seemed longer than control, pointing to a growth hastening in a low pollution levels scenario, while those exposed to the highest concentration appeared significantly shorter. This result could be interpreted as a growth retardation in highly polluted environments. This hormetic trend in tadpoles’ development has already been observed for different pesticides and heavy metals (James and Little, 2003; Smith et al., 2004; Nations et al., 2011, 2015).

In the first scenario, we can assume that the environmental stressor represented by the pollutant bursts tadpole’s development in terms of adaptive stress response, increasing length, which would eventually favour movement away from the unsuitable – polluted – medium. On the other hand, an indirect effect of low GBHs water contamination could consist in a faster development and an early metamorphosis at smaller size, with obvious negative outcomes in terms of higher predation risk and possible lower reproductive success in later-life stages (Altwegg and Reyer, 2003; Cauble and Wagner, 2005).

RU-PW caused a general growth retardation for tadpoles raised in 7.6 mg a.e./L, for both total length and maximum tail height, thus producing shorter individuals with thinner tails (Fig. 5). This could be due to the energetic cost of detoxification, reducing the amount of available energy for growing and metamorphosis. Wilkens and colleagues (2019) recently demonstrated an effect of two xenobiotics (sulfentrazone and glyphosate) and their blend on metabolic rates, oxidative stress and plasma corticosterone circulating level in tadpoles of the bullfrog *Lithobates catesbeiana*. The authors conclude that tolerance to herbicide is associated with an increase in the energy demand to keep the homeostasis and ensure the animal’s survival. A significant reduction in the embryo total length, a sensitive parameter of the teratogenesis assay in *Xenopus* (FETAX), was also observed in *Xenopus* embryos starting from the RU-PW concentration of 5mg a.e./L (Bonfanti et al., 2018).

**Figure 5:**
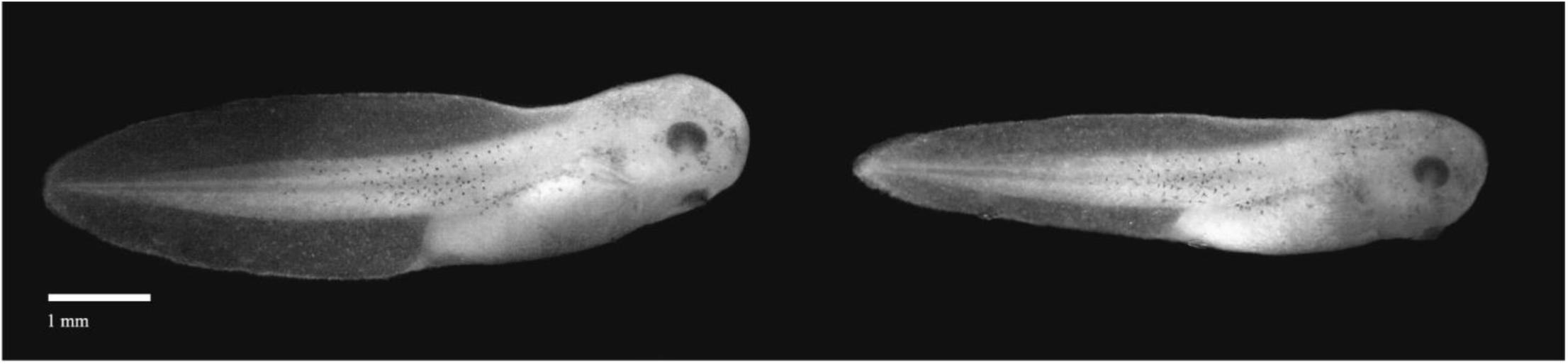
Lateral view of tadpoles after the 96h exposure from the control group (left) and RU-PW 7.6 mg a.e./L (right) showing the difference in maximum tail height.

Growth retardation may impair tadpoles’ ability to swim efficiently at hatching and therefore negatively influence their chances to cope with potential predation threats. Although in our study growth retardation was only detected at 7.6 mg a.e./L, which is the worst-case scenario concentration predicted in shallow water bodies, similar effects were found in mesocosm experiments at lower concentrations (Relyea, 2005). Nonetheless, in previous studies there seems to be differences in the effects on development and survivability of tadpoles exposed to GBHs, with lower impacts generally observed in mesocosms experiments even when testing POEA containing herbicides, the most toxic surfactant for amphibians (Mikó et al., 2015).

### Lateralization impairment at early developmental stages

Among the behavioural traits investigated in this study, the intensity of lateralization seemed to be affected by exposure to glyphosate, as significant higher values were observed for the control group when compared to the experimental group raised in 7.6 mg a.e./L RU-PW. At Gosner stage 25, we discovered that lateralization index (L_A_) was lower in exposed tadpoles compared to those raised in the control solution. Laterality is linked to anti-predatory behaviour and, therefore, an alteration caused by environmental factors could lead to a decreased efficacy of defensive responses and higher larvae mortality rates.

To date, no study has investigated how glyphosate exposure during development might affect lateralization in early vs later life stages of amphibians’ tadpoles. As Roundup® seems to activate the physiological pathway of developmental plasticity linked to anti-predator responses in tadpoles of the wood frog (*Lithobates sylvatica*), the leopard frog (*L. pipiens*) and the agile frog (*Rana dalmatina*) (Relyea, 2012; Mikó et al., 2017), and given that a higher predation risk environment during early ontogeny has been proven to lead to a higher L_A_ index in tadpoles in later life stages (Lucon-Xiccato et al., 2017), a higher L_A_ index in tadpoles treated with RU-PW during early ontogeny could have been a plausible developmental plastic response. Nonetheless, this was not the case in our experiment. RU-PW did not seem to induce behavioural and morphological adaptations linked to antipredator responses. Indeed, tadpoles raised in 7.6 mg a.e./L had a smaller tail membrane than those raised in the control solution, rather than a deeper tail membrane which is a typical morphological anti-predator change (Van Buskirk, 2001).

In the second session, when tadpoles reached Gosner stage 28/29, no significant differences between groups were detected although laterality was lower in all four groups with respect to the first session (Gosner 25; Fig. 2). So, it seems that the effects of RU-PW were only detectable shortly after hatching, and our results support the hypothesis that behavioural changes may be influenced by the developmental stages in which it falls (Mikò et al., 2017). To our knowledge, no study has yet explored how lateralization varies during tadpoles’ development and how it can be affected by environmental factors during ontogeny. If we consider the modification of the L_A_ index in the control group through development, as the natural occurring variation of the intensity of lateralization, we could hypothesize that lateralization index is higher just after hatching and then decreases at later life stages. A higher L_A_ index at earlier life stages could help tadpoles to cope with the higher early predation threats (Dadda et al., 2010), however it seems that experience is required in order to develop a certain level of behavioural lateralization (Lucon-Xiccato et al., 2017).

### RU-PW and antipredator response

The capacity of larval anurans to detect water borne cues produced by predators and properly alter their behaviour (e.g. hiding or reducing activity) is crucial for survival: tadpoles of many frog species are known to reduce level of activity after being exposed to predatory cues, in particular when chemicals are produced by a familiar predator preying upon conspecific prey (Moore et al., 2015). In our study, we did not detect any effect of the exposure during development regarding antipredator response both in the first and second trial sessions. This means that tadpoles raised in RU-PW solution had the ability to detect predatory and conspecific cues in a way that apparently resembles the ability of individuals raised in the control solution. Nonetheless, we cannot rule out that given the high concentration of predator and conspecific cue, any possible alternative responses of the distinct treatments could have been concealed by the triggering of a general strong decrease in the tadpole activity level. Another possibility is that the larvae of *P. ridibundus* are able to cope with the negative effect of the herbicide toxicity in ecological relevant concentrations, and particularly do not suffer shortcomings of the exposure at the sensory and nervous level. However, the observed differences in lateralization and basal activity (see section *Lateralization impairment at early developmental stages*) seem to support the first scenario (i.e. excess of stimulus).

### Lower basal activity in exposed tadpoles

The dramatic decrease in basal activity observed in tadpoles exposed to RU-PW, compared to those raised in the control solution in the first experimental session, may have consequences in the survivability and ability to forage of tadpoles. Indeed, foraging is crucial for survival and tadpoles need to efficiently balance the risk of starving and the risk of predation by adjusting their activity levels (Werner and Anholt, 1996). In case of general lower basal activity, tadpoles may be impaired in the foraging activity, which translates in less energy acquisition and growth and prolonged exposure to predators and water pollutants. This may result in a positive feedback cycle: higher Roundup® levels during development decreases basal activity, which may decrease foraging activity and result in lower growth rates that ultimately lead to prolonged time of exposure to water pollutants.

Bridges (1999) noted that tadpoles of *Hyla versicolor* exposed to carbaryl had significantly lower activity levels, even when no predator were present, than tadpoles not exposed to the herbicide. The consequence is a longer time spent in environments with less food availability since spending too much time resting may lower predation risk, but at the same time decreases energy acquisition.

In shallow ephemeral ponds, glyphosate concentrations are generally higher due to being located near Roundup® application and due to lack of use restrictions since they are not considered water bodies (Battaglin et al., 2009). In these environments, the reduced growth rate caused by the herbicide may increase the risk of not achieving metamorphosis before the water dries out (Bridges, 1999). Furthermore, as water stratification does not occur in shallow water bodies, a phenomenon which causes glyphosate and surfactants to concentrate near the surface, the risk of exposure increases even more for the offspring of amphibians that breed in ephemeral ponds (Jones et al., 2010).

At Gosner stage 28/29, we observed a reduction in the basal activity of tadpoles from the control group, but no significant variation was observed among all the treatments. Despite this significant drop in activity for the control group, tadpoles raised in 7.6 mg a.e./L still showed similar activity levels to the control group in the second session. One possible explanation for the observed shift in the basal activity of tadpoles exposed to RU-PW is that the exposed tadpoles may have increased their activity to compensate the initial negative effect of the herbicide. Alternatively, the control group may have decreased its basal activity to optimize their growth and to storage the required amount of energy for further metamorphosis.

## CONCLUSIONS

Amphibians are experiencing a decline on a global scale, mainly due to human activity (Carey and Bryant, 1995; Stuart et al., 2008). One of the main factors considered to negatively impact amphibians’ conservation is the environmental pollution through pesticides employment, which are often directly applied to the soil and may contaminate the aquatic environment through leaching, wind or transported by runoff waters (Collins and Storfer, 2003; Saunders and Pezeshki, 2015).

Our study demonstrates that tadpoles of *P. ridibundus* are sensitive to a glyphosate-based herbicide in terms of both morphological and behavioural modifications. Tadpoles exposed developed faster at low concentrations (0.7 and 3.1 mg a.e./L), while were affected by overall growth retardation at the highest concentration tested (7.6 mg a.e./L). Lateralization and basal activity were affected by embryonic exposure to glyphosate, however we could not detect any effect on antipredator behavioural responses. Some of the observed modifications may be attributed to altered or reduced brain development, or to the inflammation and consequent infiltration of eosinophilic granule cells/mast cells in neuronal bodies, as demonstrated by previous laboratory tests involving both non-model or model organisms (Ramírez-Duarte et al., 2008; Bonfanti et al., 2018). Although our study reports the results of acute exposure to glyphosate, it can be assumed that, after prolonged exposure, the observed behavioural alterations would only worsen, making tadpoles less responsive to stressful synergetic situations such as habitat fragmentation, UV radiation, pollutants, pathogenic agents, invasive species, and predators. All these factors, among others, would contribute to the decrease of populations of this and other similar species.

To the best of our knowledge, this is the first study describing variation in lateralization in tadpoles of the genus *Pelophylax*, which comprise nearly 20 taxa spread throughout Europe and Asia. Noteworthy, *P. ridibundus* has been widely translocated among European countries, and it is known to hybridize with local taxa with unknown ecological consequences. In Northern Italy, where this species has undergone multiple release, hybridization with *P. lessonae* (the pool frog) and particularly *P. esculentus* (the edible frog, which carries the genome of the latter two species as a result of hybridogenetic mechanisms; Berger, 1973), may eventually impact the species-specific response of native species to environmental pressures like predation in polluted environments. *P. ridibundus* is overall less sensitive to disturbance than the native taxa, particularly *P. lessonae*, therefore it can be assumed that observed effects of herbicide would be more evident in less tolerant tadpoles.

Nonetheless, the latter statement needs to be supported by further studies, in order to clarify if marsh frogs may be further advantaged on native taxa via pollutant resistance.

Finally, since aquatic environments are essential to both the life cycle of amphibians and their reproductive success, in this study we supported the hypothesis that water contamination may greatly impair the survivability of amphibian populations.

## Acknowledgements

We thank F. Pupin and A. Bianchi for helping in fieldwork activity, and S. Seghizzi for her support in laboratory analysis. Permission for frog collection and suppression were granted to A. Bellati by the Italian Ministry of the Environment under Prot. N. 0003221.15-02-2017.

